# A site specific model and analysis of the neutral somatic mutation rate in whole-genome cancer data

**DOI:** 10.1101/122879

**Authors:** Johanna Bertl, Qianyun Guo, Malene Juul, Søren Besenbacher, Morten Muhlig Nielsen, Henrik Hornshøj, Jakob Skou Pedersen, Asger Hobolth

## Abstract

**Background:** Detailed modelling of the neutral mutational process in cancer cells is crucial for identifying driver mutations and understanding the mutational mechanisms that act during cancer development. The neutral mutational process is very complex: whole-genome analyses have revealed that the mutation rate differs between cancer types, between patients and along the genome depending on the genetic and epigenetic context. Therefore, methods that predict the number of different types of mutations in regions or specific genomic elements must consider local genomic explanatory variables. A major drawback of most methods is the need to average the explanatory variables across the entire region or genomic element. This procedure is particularly problematic if the explanatory variable varies dramatically in the element under consideration.

**Results:** To take into account the fine scale of the explanatory variables, we model the probabilities of different types of mutations for each position in the genome by multinomial logistic regression. We analyse 505 cancer genomes from 14 different cancer types and compare the performance in predicting mutation rate for both regional based models and site-specific models. We show that for 1000 randomly selected genomic positions, the site-specific model predicts the mutation rate much better than regional based models. We use a forward selection procedure to identify the most important explanatory variables. The procedure identifies site-specific conservation (phyloP), replication timing, and expression level as the best predictors for the mutation rate. Finally, our model confirms and quantifies certain well-known mutational signatures.

**Conclusion:** We find that our site-specific multinomial regression model outperforms the regional based models. The possibility of including genomic variables on different scales and patient specific variables makes it a versatile framework for studying different mutational mechanisms. Our model can serve as the neutral null model for the mutational process; regions that deviate from the null model are candidates for elements that drive cancer development.

## 1 Background

Cancer is driven by somatic mutations that convey a selective advantage to the cell. However, in most cases the somatic mutation rate in cancer cells is considerably higher than in healthy tissues, of which only a small fraction of the mutations are thought to be associated with cancer development [Stratton et al., 2009]. The majority of the mutations are neutral and is caused by perturbed cell division, maintenance and repair or over-expression of mutagenic proteins (e.g. the APOBEC gene family [Bacolla et al., 2014]).

A comprehensive framework of the random mutation process in cancer cells is key to identify the regions, pathways and functional units that are under positive selection during cancer development. The mutation rate varies along the genome, depending on genomic properties of the position such as the sequence context (e. g. the 5’ and 3’ nucleotides; [Alexandrov et al., 2013]), chromatin organisation [Polak et al., 2015] or replication timing [Lawrence et al., 2013]. Many studies have investigated what determines the mutation rate and what kind of models should be used. Lawrence et al. [2013] modeled the mutation heterogeneity using local regression with expression level and replication timing as explanatory variables. Melton et al. [2015] implemented a Poisson-binomial model on 50 kb windows using the annotation of the transcription factor binding site and replication timing. Polak et al. [2015] applied random forest regression on mutation counts in 1Mb windows using histone modification and density of DNase I hypersensitive sites. Lochovsky et al. [2015] predicted the number of mutations per element by a beta-binomial distribution using replication timing and noncoding annotations such as promoter, UTR and ultra-conserved sites. Unlike these publications that segmented the genome into regions according to the explanatory variables and estimated separate models for different regions, we propose a framework based on site-specific multinomial regression where this division is not necessary. We can use site-specific explanatory variables without dividing the data into subsets. In this way, we use the full dataset to estimate the regression coefficients for all the explanatory variables. We further include GC content and CpG island annotations to describe the local context composition for genomic regions. We include the conservation score phyloP to study the correlation between somatic mutations and germline substitutions. We also include an explanatory variable to mask the repeat regions in the genome as due to technical reasons, mutation calls at repeat regions are biased.

Generalized linear models also provide an interpretable framework for modeling and hypothesis testing. For example, we can estimate the mutation rates of CpG sites within and outside CpG islands, and test if they are different, and a patient-specific intercept allows us to take the large variation of mutation rates between patients into account.

Here, we analyse 505 cancer genomes from 14 different cancer types [Fredriksson et al., 2014]. We compare the performance in predicting mutation probabilities of region-based Poisson models, site-specific binomial models and site-specific multinomial models. The site-specific multinomial model can predict both the overall mutation rate and mutation types accurately. We use a forward model selection procedure to compare and identify the explanatory variables that best explain the heterogeneity of the site-specific mutational process (Figure 1).

**Figure 1:**
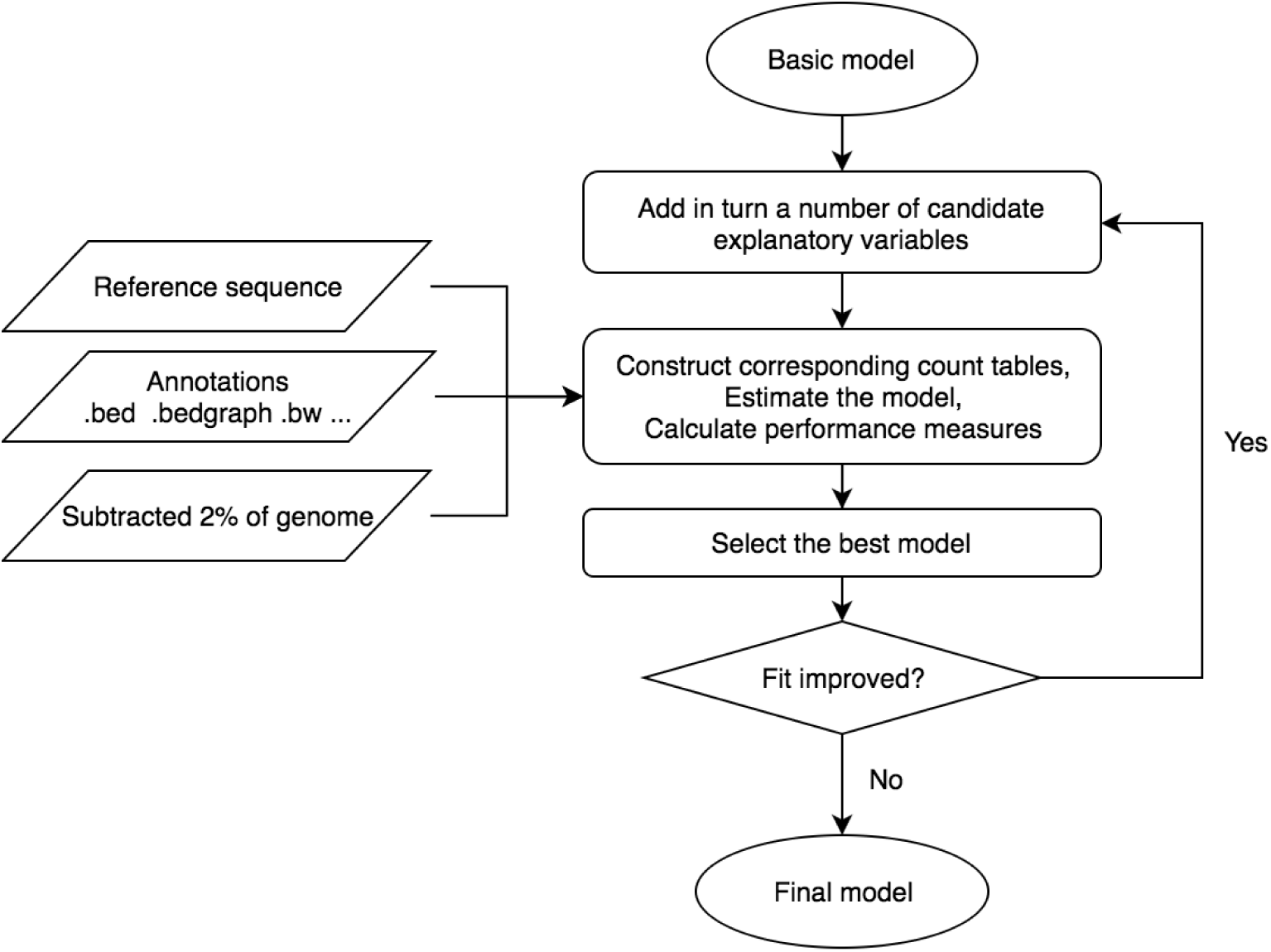
Workflow of the forward model selection procedure. The forward model selection is implemented on 2% of the data to determine the explanatory variables included in the final model. In each iteration of the model selection, data tables are generated to summarize the site-specific annotations. The performance of the models are measured with McFadden’s pseudo *R*^2^ and deviance loss obtained by cross-validation. The explanatory variable with the best performance is included in the set of variables for the next iteration. Parameter estimation for the final model is based on the remaining 98% of the data.

The forward model selection procedure is implemented using 2% of the data while the final model fit is obtained from the remaining 98%. We find that site-specific conservation (phyloP), replication timing and expression level are the best predictors for the mutation rate. In general, the framework allows formal testing for inclusion of explanatory variables and also interaction terms. The impact of different explanatory variables can be inferred from the parameter estimates of our final model as the multiplicative changes in mutation rate. Our analysis confirms some known mutational signatures and it identifies associations with genomic variables like replication timing. It can also be used as the null model for cancer driver detection [Juul et al., 2017] and other applications that rely on a model of the mutation rate (e. g. identification of the tissue of origin for tumors of unknown primary, Polak et al. 2015).

## 2 Results

### 2.1 Heterogeneity of the mutation rate

We observe significant heterogeneities of the mutation rate at multiple levels (Figure 2). The mutation rate varies among cancer types (Figure 2a): skin cutaneous melanoma, colorectal cancer and lung adenocarcinoma are the cancer types with the highest mean mutation rates (5 – 10 mut/patient/Mb) while thyroid carcinoma, prostate adenocarcinoma and low-grade glioma are the lowest (0.5 – 1 mut/patient/Mb). The mutation rate also differs between samples from the same cancer type, with the largest variation seen for skin cutaneous melanoma (the mutation rate ranges from 1 mut/patient/Mb to 150 mut/patient/Mb).

**Figure 2:**
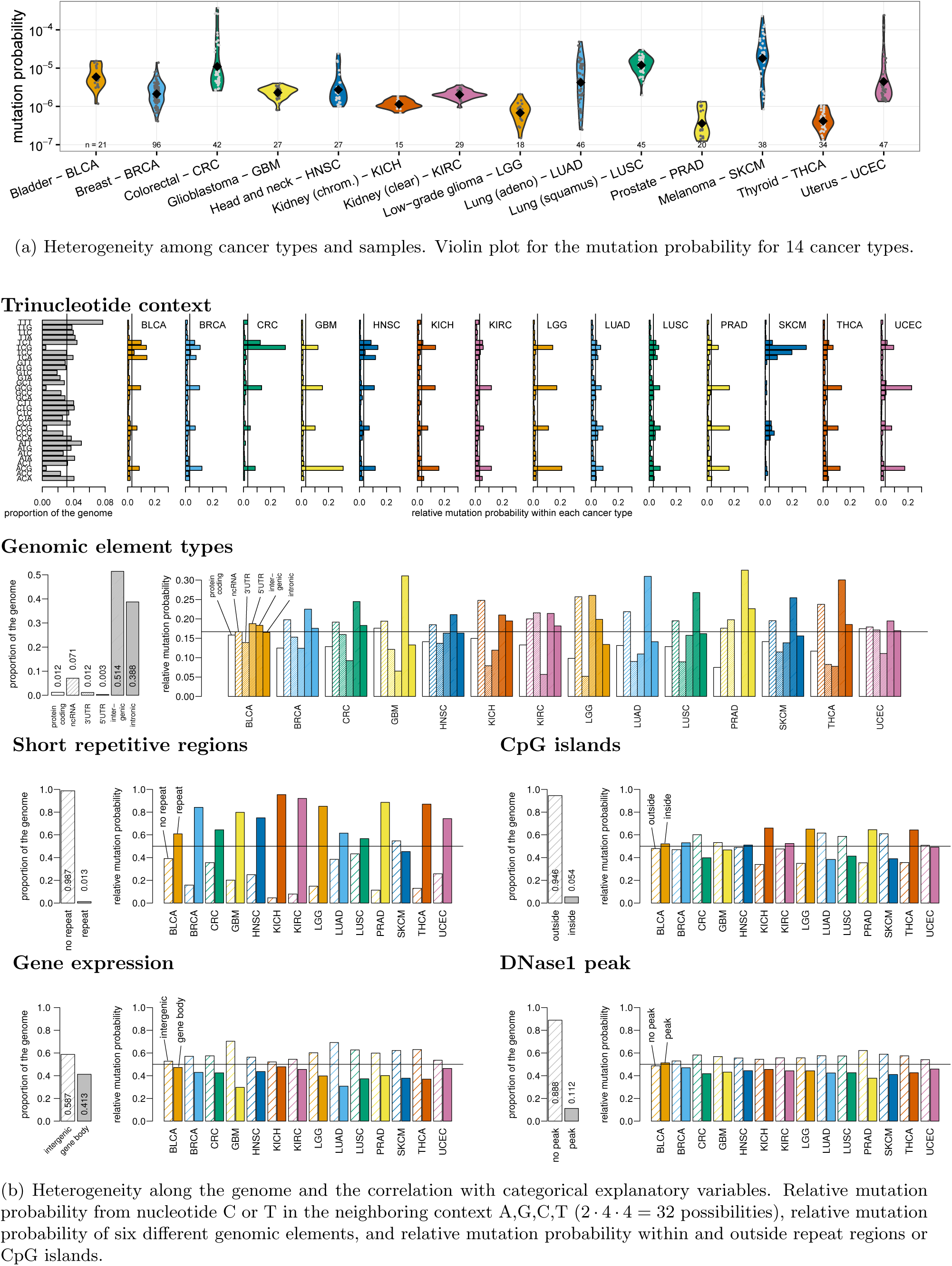
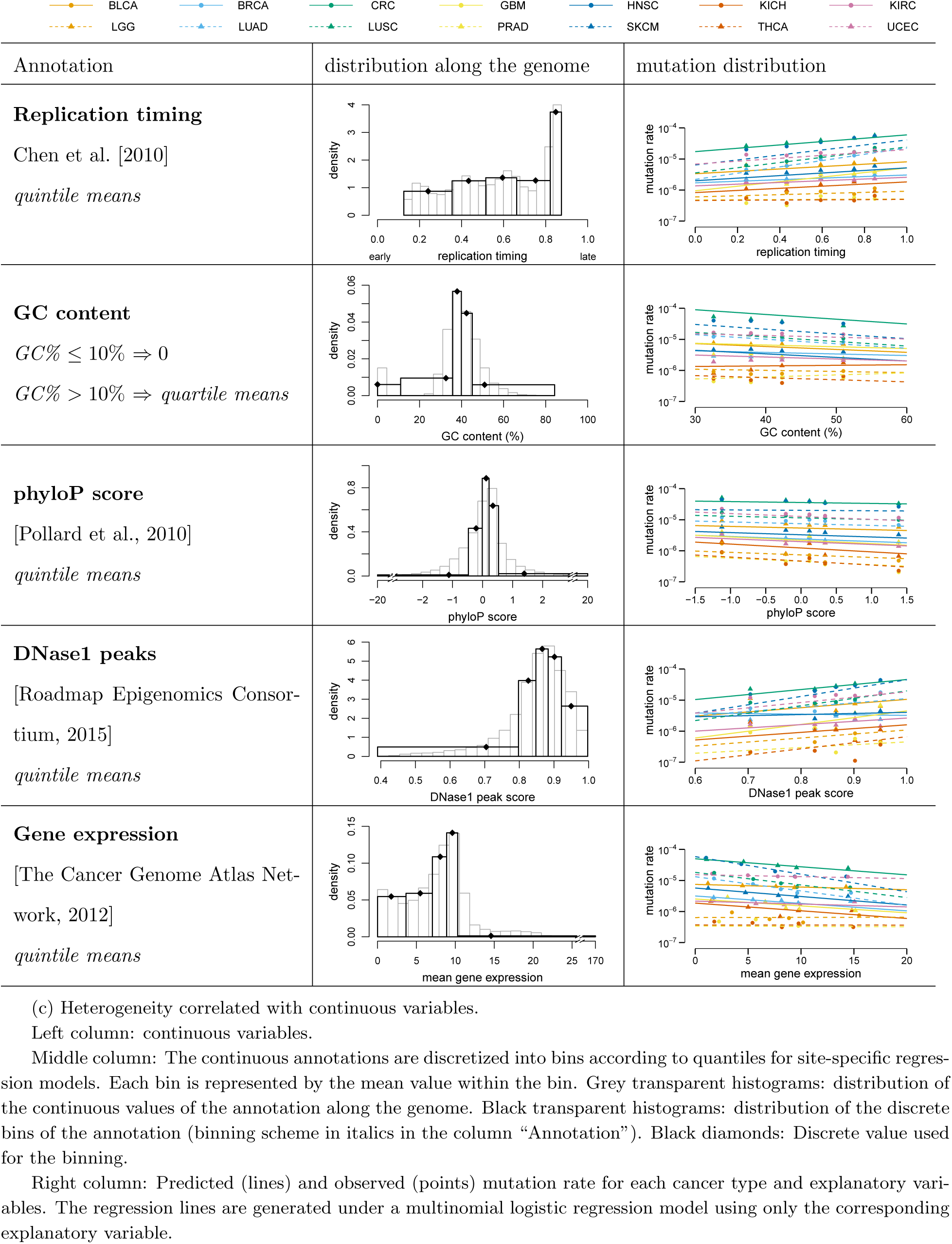
Heterogeneity of the mutation rate and explanatory variables (continues on the next page).

The mutation rate also varies between different genomic contexts (Figure 2b). As previously shown, we find that mutational signatures are cancer type specific by looking into the mutation rate for different trinucleotide contexts. The mutation rate at TT sites is higher in skin cutaneous melanoma than in any other cancer types, and the mutation rates at CpG sites are elevated for all the cancer types. In colorectal cancer, we observe more mutations at TCG sites and in glioblastoma more mutations at ACG sites.

The mutation rate also differes between genomic environments defined by the explanatory variables (Figure 2b, 2c). Coding regions tend to have fewer mutations for all cancer types. Mutation rates are elevated for simple repeat regions, which might be related to mapping artefacts and ensuing technical challenges during mutation calling. The effect of CpG islands varies for different cancer types. The mutation rate in CpG islands is higher than in regions outside for thyroid carcinoma, prostate adenocarcinoma, low-grade glioma and kidney chromophobe, while for colorectal cancer, lung adenocarcinoma, lung squamous cell carcinoma and skin cutaneous melanoma the situation is reversed. Regions that are late replicated, GC rich, evolutionarily less conserved, inside DNase1 peaks and lowly expressed have an elevated mutation rate. The explanatory power of the variables varies across cancer types as shown by the regression lines in Figure 2c.

### 2.2 Granularity of regression models

We model the mutation probability in cancer genomes using a set of regression models. The most coarse-grained description of the number of mutations in a region is a Poisson regression count model and the most fine-grained is a binomial or multinomial site-specific regression model. Here we describe and investigate in detail the (dis)advantages of these three models. A conceptual overview of the models is given in Figure 3a.

**Figure 3:**
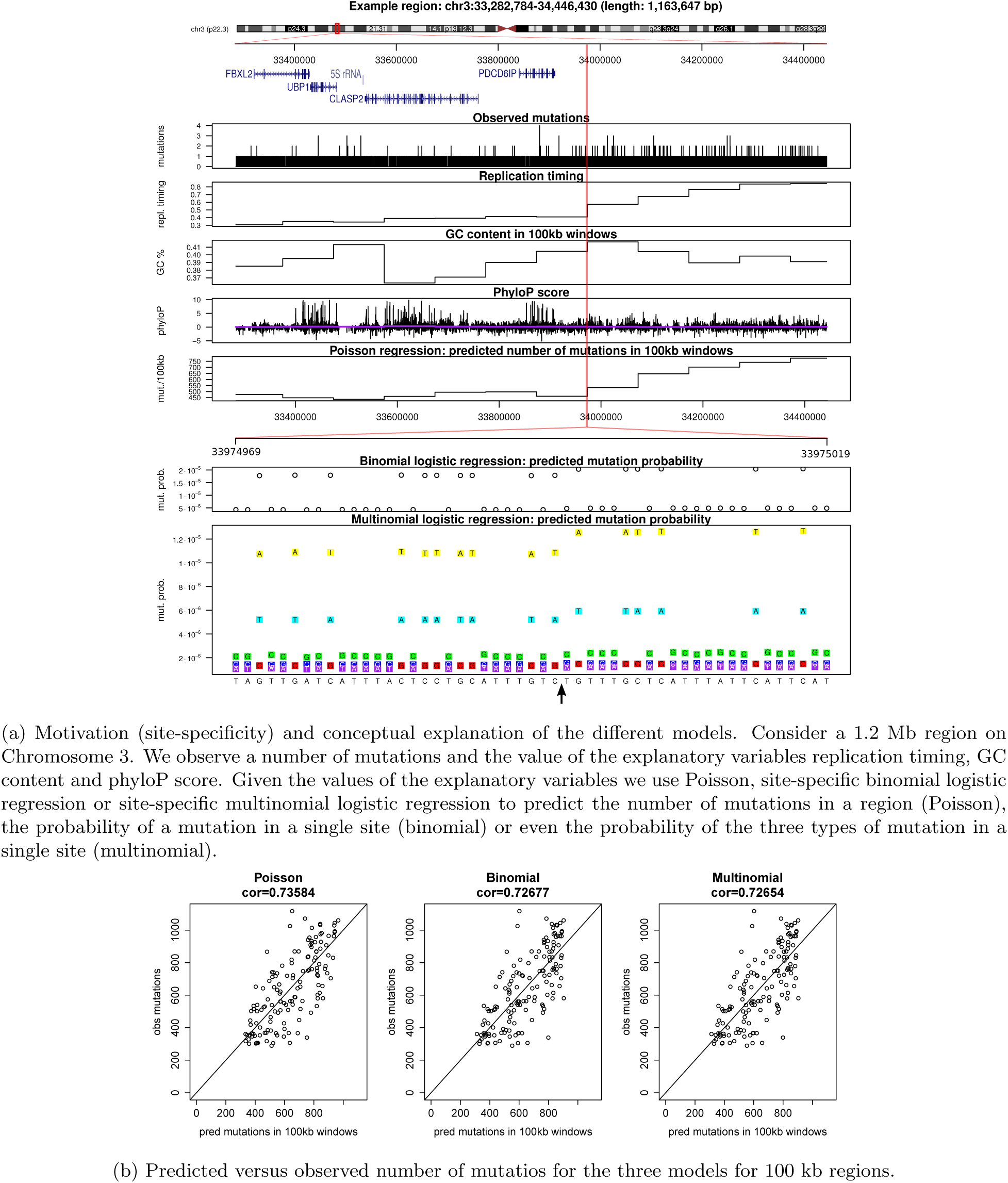
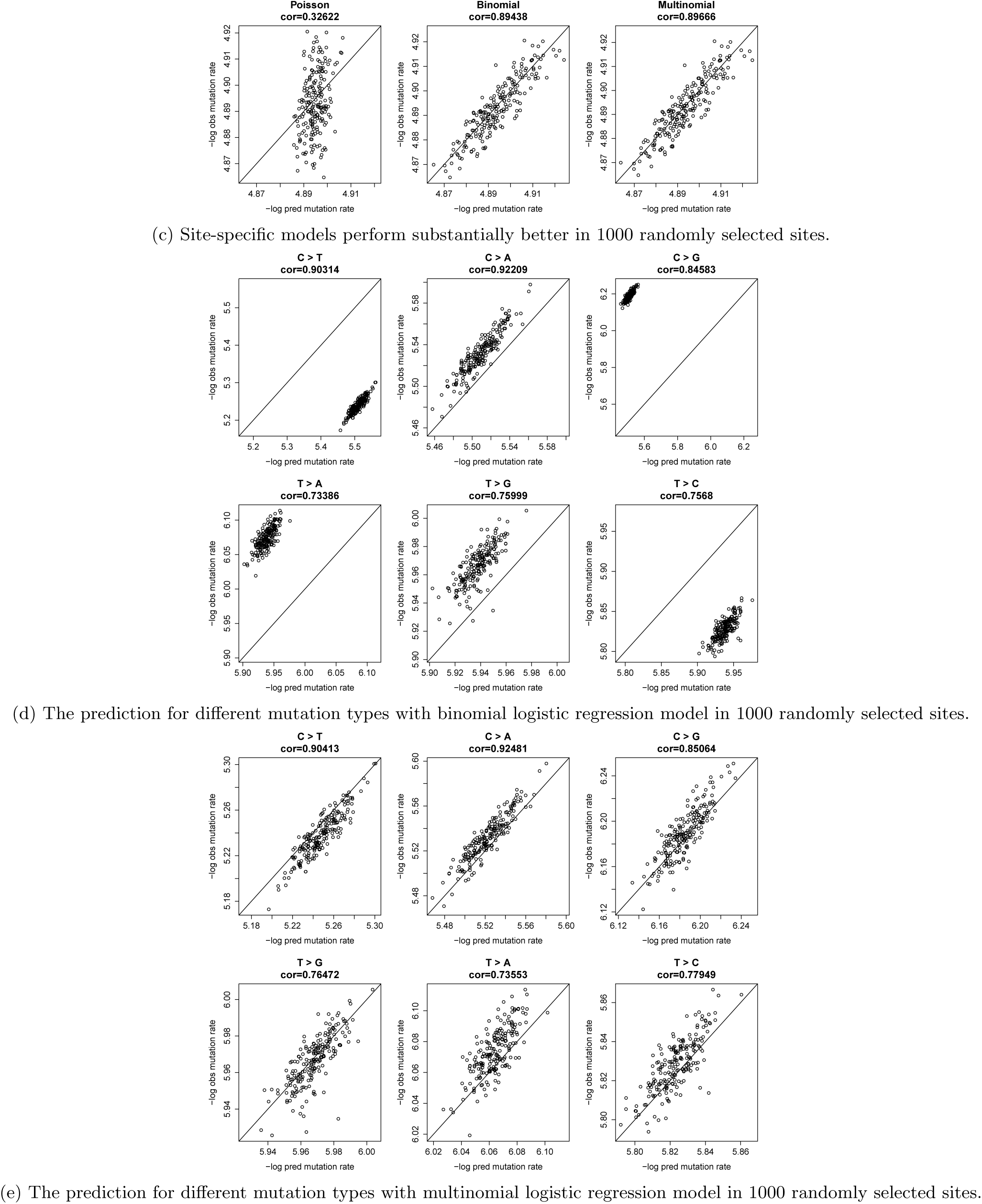
Comparison of Poisson regression model, site-specific binomial logistic regression model and site-specific multinomial logistic regression model (continues on the next page).

#### Poisson count regression model

In Poisson regression, the number of mutations in a genomic region of fixed length is modeled. The whole genome is divided into regions of pre-fixed length or according to the value of explanatory variables (e.g. segmented by genomic element types). Regression modeling is facilitated by summing the mutation counts and summarizing the annotations over the region.

We model the mutation count *N_r_*_,*sam*_ in the *r*-th genomic region with length *L_r_* in sample *sam.* Furthermore, *can* is the cancer type of the sample. Mutations arise randomly with probability *p_r_*_,*sam*_. As *p_r_*_,*sam*_ is small and *L_r_* is large, we have the Poisson approximation of the binomially distributed number of mutations

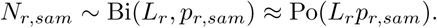

For *J* explanatory variables, the expected mutation count *λ_r_*_,*sam*_ = *L_r_p_r_*_,*sam*_ can be modeled by a Poisson regression:

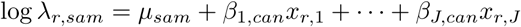

where for the *j*th explanatory variable, *x_r_*_,*j*_ is the average value of the annotation across region *r* if the explanatory variable is continuous and for categorical explanatory variables, *x_r_*_,*j*_ is derived from the proportions of different levels of the annotations in region *r, j* = 1, … *, J.*

#### Site-specific binomial regression model

In site-specific regression models, the mutation probability is modeled in each position of the genome. We enable regression modeling by binning the continuous annotations, such that we are able to sum mutation counts over positions with the same combination of annotations, and thereby reduce the size of the data set. We consider both site-specific binomial and multinomial regression models.

We model the mutation probability *p_i_*_,*sam*_ at a site *i* in sample *sam* of cancer type *can.* With a logit link, the mutation probability can be modeled by logistic regression:

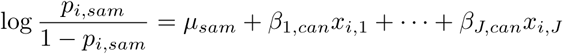

where *x_i_*_,*j*_ is the value of the *j*th explanatory variable at site *i.*

#### Site-specific multinomial regression model for strand-symmetric mutation types

We model the mutation probability for different mutation types. Assuming strand-symmetry, we are not distinguishing between e.g. A>G (A to T) mutations and T>C mutations, *p^A^*^>*G*^ = *p^T^*^>*C*^. We consider the strand with the C or T nucleotide, and the mutation probability matrix is

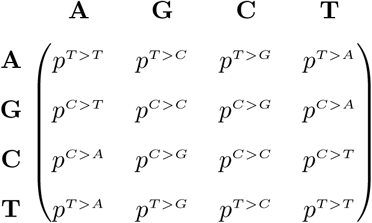

with only 6 types of mutations.

We model these mutation probabilities by setting up two independent multinomial logistic regression models, one for mutations from (G:C) base pairs and one for mutations from (A:T) base pairs. The (G:C) basepairs are modelled with probabilities

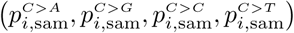

for (G:C) position *i* in sample *sam.* With *J* explanatory variables, the mutation probability at G:C position *i* in sample *sam* of cancer type *can* can be written as:

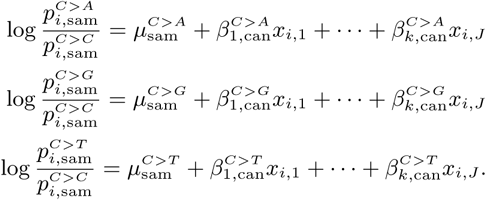

Note that the probability for no mutation 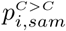 is the reference. Similarly, (A:T) basepairs are modeled with probabilities

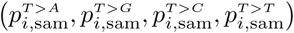

and the reference is the probability for no mutation 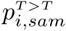. We compare the performance of the three models on 2% of the whole genome data. The setting for the three models is shown in Figure 3a. Each model is trained with replication timing, phyloP, and context information from the reference genome. For the region-based Poisson model, continuous annotation values are averaged over the selected region and GC percentages are calculated for each region. For the site-specific models, we use the site-specific annotations for each site. Continuous values are discretized to simplify the estimation process.

The results are shown in Figure 3b. When predicting mutation counts in large windows (100 kb), the three regression models perform similarly. For prediction in a randomly selected small number of sites (1 kb), the two site-specific models out-perform the region-based model (Figure 3c). The site-specific models can capture the mutational heterogeneities between sites and provide a more accurate mutation probability at any resolution. This is in contrast to the region-based model where a large number of sites are required for accurate predictions. In addition to predicting the probability for mutation events, the multinomial regression model can also predict the mutation probabilities for different mutation types (Figure 3d, 3e).

### 2.3 Model selection

We consider the site-specific multinomial regression model to predict mutation probabilities for different mutation types at a single site. We implement a forward model selection procedure to determine the explanatory variables in the final model (Figure 1). In each step, we add all possible new variables to the previous model in turn and rank the resulting new models. We identify and include the explanatory variable with the best fit and iterate the procedure several times. Cross-validation is used to assess the fit of a model using the deviance loss function while avoiding overfitting (Section 4.3.1). By construction, the fit improves during the model selection procedure (Figure 4a). We also evaluate McFadden’s pseudo *R*^2^ as a measure of the explained variance that is valid in categorical regression models (Section 4.2.3).

**Figure 4:**
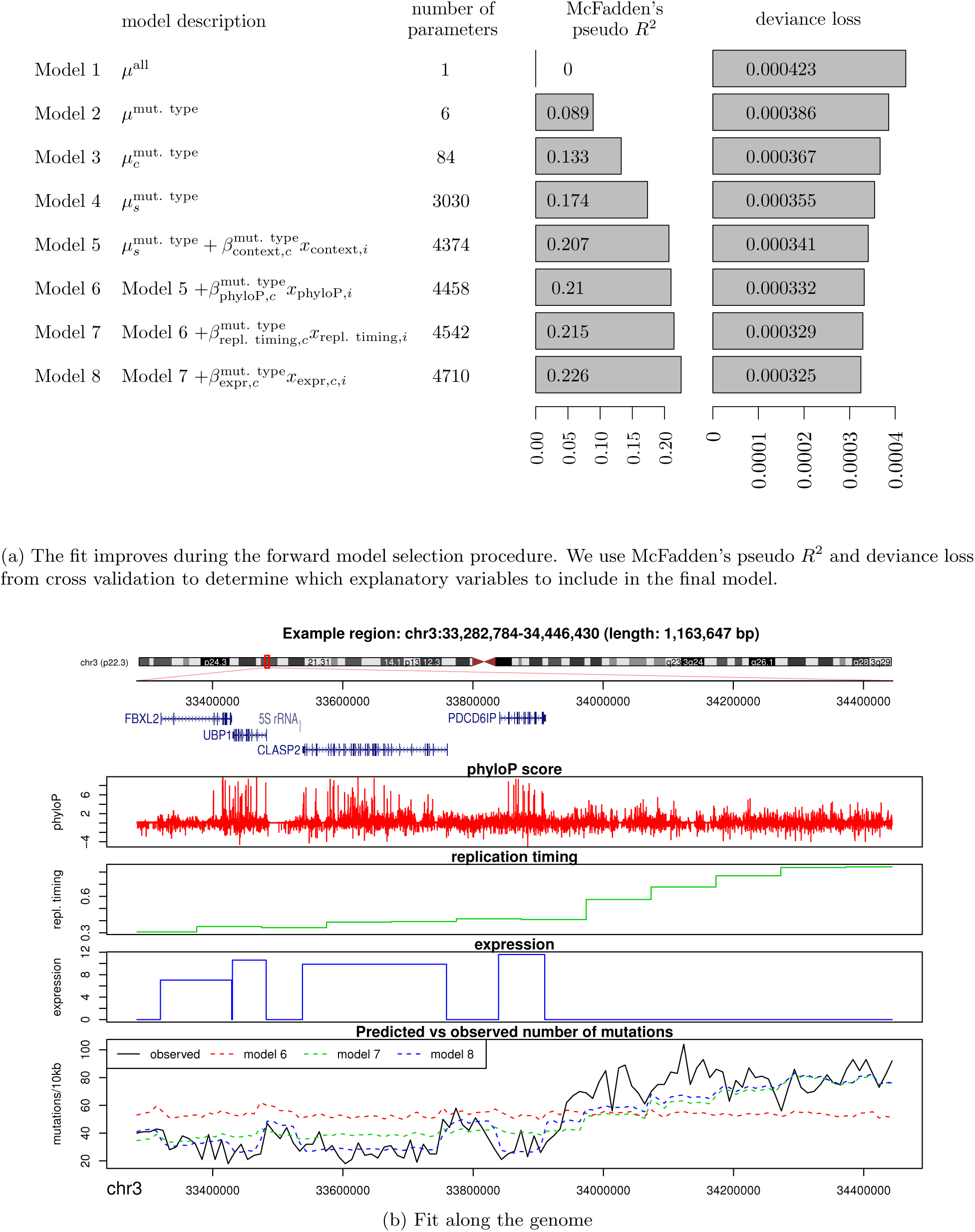
Model selection results. Note that the observed number of mutations is better predicted for the more complex models.

As a reference model, we start out with a single mutation rate for the whole genome in all samples (Model 1; Figure 4a). This model cannot explain any of the variation in the mutation rate between samples and positions, so McFadden’s pseudo *R*^2^ = 0. After including the six strand-symmetric mutation types in the model (Model 2), we add cancer and sample specific intercepts (Model 3 and Model 4) to make sure that we account for sample-specific mutation rates. In the next step, we include the left and right neighboring base-pair for each cancer type (Model 5).

Starting with Model 5, additional annotations are added using forward model selection procedure. We consider the phyloP score, replication timing, expression, genomic segments, GC content in 1 kb, CpG islands, simple repeats and DNase1 hypersensitivity. For each of these variables, cancer specific regression coefficients are estimated to allow for differences in the mutational process between cancer types. For expression, we use data directly obtained from matching tumor types, so we also take expression differences between cancer types into account.

The annotation with the largest decrease in the deviance loss function is the phyloP score (Model 6). Subsequently, replication timing (Model 7) and gene expression (Model 8) are added. At this point, we stop the model selection procedure to avoid the time-consuming creation of larger count tables (see Section 4.1.2 for details) as we also see that the improvement of the fit levels off. Detailed results for each step of the forward selection procedure are provided in the Appendix.

While the trinucleotide context and the phyloP score vary on a basepair scale, both replication timing and expression vary on a kilo-base scale. Even though much of the per-base-pair variation in the mutation rate is already captured in Models 5 and 6, the long-range variation is considerably better explained in Model 7 and Model 8, as illustrated in Figure 4b.

We can see that adding replication timing in Model 7 considerably changes the predicted mutation rate obtained from Model 6 that is relatively uniform in the 10 kb scale shown in the figure, by taking the replication timing gradient in the region into account. Finally, in Model 8, the lower mutation rate in highly expressed regions is added, which again improves the prediction.

### 2.4 Estimation results

Upon determining the final model from the model selection procedure, we estimate parameters for the multinomial logistic regression model on the remaining 98% of the genome. To study the difference between cancer types, all position-specific explanatory variables are stratified by cancer type. The coefficients represent multiplicative changes in mutation rate. Our results confirm the large differences in mutation pattern both between samples and cancer types, but also between different genomic and epigenomic regions.

We observe that regions that have been highly conserved during human evolution have a lower mutation rate in all cancer types and for all mutation types, and this difference is nearly always significant (Figure 5a). For kidney chromophobe and prostate adenocarcinoma, it is reduced to less than half for some of the mutation types. In breast cancer, head and neck squamous cell carcinoma, kidney chromophobe and thyroid carcinoma, this difference is significantly more pronounced at A:T positions than at G:C positions, but there is no general pattern with respect to the mutation type.

**Figure 5:**
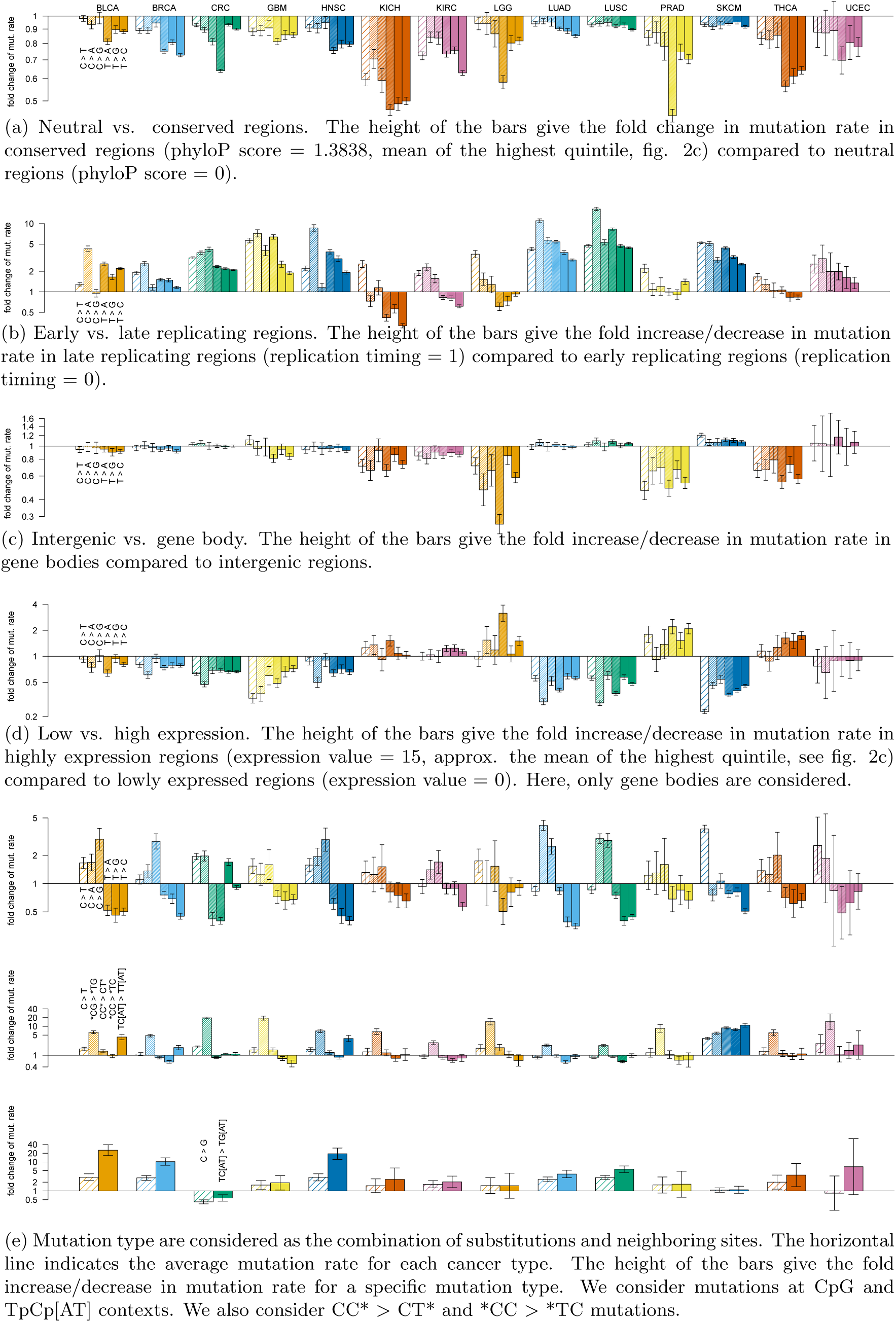
Parameter estimation results.

As previously described [Lawrence et al., 2013], we nearly always find a positive association between replication timing and mutation rates (see regression lines in Figure 2c; later replicating regions have more mutations), but the regression coefficient varies significantly between the different cancer types and the mutation types (Figure 5b). This can either point to different mutational mechanisms, or it shows that the replication timing dataset at use (Chen et al., 2010; measured on HeLa cell lines) is not equally representative for different types of cancer tissues.

For the dummy variable that distinguishes gene bodies, where expression is measured, from intergenic regions, we see a mixed pattern of insignificant and negative regression coefficients where the mutation rate in gene bodies is reduced to up to one third of the rate in intergenic regions (low-grade glioma; Figure 5c). The only exception is melanoma with a slightly increased mutation rate in gene bodies.

Also when we consider regions within gene bodies, we find a reduced mutation probability in highly expressed regions for most cancer types. The most extreme example is the probability of a C > T mutation in highly expressed regions in melanoma, which is only one fifth of the probability in lowly expressed ones (Figure 5d). Low-grade glioma, prostate adenocarcinoma and thyroid cancer show the opposite pattern for some of the mutation types, though. However, this effect is dampened by the dummy variable for the gene bodies that have comparably large coefficients of the opposite sign for these three cancer types.

We find that the mutation rates differ between the different types of mutations, but also between the specific contexts that we consider: CpG, to capture the pattern of spontaneous deamination, and TpCp[AT], to capture the APOBEC signature. We find that the C > T mutation rate is higher in CpG sites than in other sites in all cancer types. In skin cutaneous melanoma, we also observe elevated mutation rate for CC context, which is related to the elevated CC > TT mutation rate due to UV light signature [Rass and Reichrath, 2008]. We observe elevated rates of mutations that fit the APOBEC pattern in breast cancer bladder urothelial carcinoma, head and neck squamous cell carcinoma, lung squamous cell carcinoma and skin cutaneous melanoma.

## 3 Discussion and Conclusion

We use a multinomial logistic regression model to analyse the somatic mutation rate in each position of a cancer genome. We consider various genomic features, such as the local base composition, the functional impact of a region and replication timing. Because of the site-specific formulation, the model is the most fine-grained description of the mutation rate, while it can still take genomic properties into account that vary or are measured on a longer scale, like replication timing and expression levels.

The mutational spectrum and intensity are known to vary considerably between cancer types as well as between patients with the same cancer type [Lawrence et al., 2013]. In our analyses, we capture a considerable part of the variance by including cancer type and sample specific mutation rates.

We found the site-specific phyloP conservation scores to explain more than any of the other explanatory variables we tested. The conservation scores reflect the germline substitution rate between species through evolution and are designed to capture the effect of selection [Pollard et al., 2010]. However, somatic evolution of the cancer genome is not subject to the same constraints as germline evolution. In particular, purifying selection may be relaxed as many genes and regulatory elements that are needed during the organismal life cycle are dispensable for the growth of cancer cells [De Silva et al., 2014]. Instead, the conservation scores’ explanatory power in our study may stem from their correlation with genes and other functional elements, with special properties that affect their mutation rate. In particular, it is known that the mutation rate is elevated at transcription factor binding sites in some cancer types [Sabarinathan et al., 2016]. Furthermore, the phyloP scores are also correlated with expression regions and open chromatin, which both have decreased mutation rates due to transcription-coupled repair and other repair mechanisms ([Su et al., 2015]).

The second explanatory variable added is replication timing. This has not only been found for cancer genomes [Lawrence et al., 2013], but also for germline mutations [Stamatoyannopoulos et al., 2009] and somatic mutations in healthy tissue [Koren et al., 2012]. In late replicating regions, single stranded DNA (ssDNA), which is susceptible to mutagenic processes like deamination, accumulates [Stamatoyannopoulos et al., 2009]. Varying intensities of such mutational processes could explain the differences in the impact of replication timing between the cancer types and the mutation types. The different strengths of the associations between replication timing and the mutation rate can also be attributed to differences in replication timing between cell types [Hansen et al., 2010] that may be missed in the replication timing dataset at use, which is obtained from HeLa cell lines [Chen et al., 2010].

Only after the phyloP score and replication timing, the cancer type specific expression level is added to the model. This can be explained by the fact that the phyloP score and expression are correlated, as mentioned above, and so are replication timing and expression [Rhind and Gilbert, 2013]: protein-coding genes are concentrated in early replicating regions. However, expression levels that we use are tissue specific and might therefore improve the fit of the model further for that reason.

By separately estimating a regression coefficient for the gene-body and one for the expression level within gene bodies, we can potentially distinguish the general properties of genes, like location with respect to replication timing origins and sequence composition, and the intensity of transcription-coupled repair (TCR). Since TCR is a subpathway of nucleotide excision repair (NER), it is expected to act on helix-distorting mutations like for example the well-known UV light induced CC > TT mutations in melanoma. Thus, the cancer specific differences that we see might be explained by varying effectiveness of TCR, but also by different proportions of mutations that create bulky distortions. For example, the rate of C > T mutations is further decreased in highly expressed regions in melanoma than the other mutation types.

The parameters can be interpreted as multiplicative mutation rate changes. By using interaction terms, it is straightforward to analyse and test differences in mutation rate between cancer types, samples or specific genomic regions of interest. Furthermore, generalized linear models come with a natural framework for hypothesis testing. It would be interesting to compare our model-based description of the mutation rate to unsupervised learning of mutational signatures by matrix factorization [Alexandrov et al., 2013].

In the logistic regression model, we can estimate cancer type specific regression coefficients to capture the differences between cancer types. We can also include tissue specific explanatory variables if there are measurements for the corresponding cancer tissue or matching healthy tissue available. This is of particular interest for epigentic measurements like histone modifications, where the tissue of origin has been shown to be informative [Polak et al., 2015].

Patient-specific characteristics can also be added as explanatory variables. The age of the patients could be used to study clock-like mutational processes [Alexandrov et al., 2015]. Known somatic or germline mutations that are associated with specific mutational processes or repair pathways can also be used as explanatory variables, e. g. a germline deletion of *APOBEC3B* that fuses APOBEC3A with the 3’ UTR of APOBEC3B has been found to be associated with an increased number of APOBEC-type mutations [Nik-Zainal et al., 2014]. The impact of this mutation could be studied by including an interaction term with the TpCp[AT] positions.

Our model is very flexible and versatile and the explanatory variables can be customized according to different applications. An important application of the model is the prediction of the somatic mutation probability under the assumption of neutrality. In our driver detection method ncdDetect [Juul et al., 2017], we use the position-specific predictions that we obtain from the multinomial model to evaluate if the mutation rates or their distributions are significantly different from the expectation under neutrality. This allows a flexible analysis of regions of any size. Even non-contiguous regions with very different properties than the overall genomic patterns can be investigated.

## 4 Methods

### 4.1 Data

#### 4.1.1 Somatic mutation dataset

We use SNV calls from 505 tumor-normal samples across 14 different cancer types from Fredriksson et al. [2014]. We build our data set based on the UCSC hg19 assembly. We removed regions with low mappability, ultra-high mutation rates and lacking annotation. Problematic regions for NGS alignments identified for the ENCODE project [The ENCODE Project Consortium, 2012] were subtracted. Low mappability regions, defined by the GEM tool [Derrien et al., 2012] and CRG Alignability track from UCSC with mappability less than 0.5 in 100-mers, were also subtracted. Hyper-mutated genomic segments containing Immunoglobulin/Tcell receptor (IG/TR) genes defined by GENCODE together with 10 kb flanking regions, combined when less than 100 kb in distance, were excluded from analysis. We also excluded sites on ChrX and ChrY, because some of the annotation files we use lack information for one or both of the sex chromosomes.

A total number of 14, 200, 393 SNVs are observed in the subtracted regions for 505 samples across 14 cancer types.

##### Genomic element types

We divided the genome into six genomic annotations: coding, 5’ UTR, 3’UTR, ncRNA, intron and intergenic. Based on the GENCODE v.19 transcripts, coding regions, 5’ UTR regions and 3’ UTR regions as well as introns were defined for protein-coding transcripts. Non-coding RNA regions were defined as all remaining regions in the full transcript set. All remaining bases were categorized as intergenic.

##### GC content

We calculate the percentage of G:C base pairs (strong sites) in 1 kb windows based on the reference genome. Regions with GC percentage less than 10% are annotated with value 0. Other regions are discretized into quartiles.

##### CpG islands

We segmented the genome by presence or absence of CpG islands. The CGI Mountain annotation from CgiHunter (http://cgihunter.bioinf.mpi-inf.mpg.de/) was used. The CGI Mountain score quantifies if a region is a CpG island. Scores above zero indicate a CpG island. We use a dummy variable derived from the CGI Mountain score in our analysis indicating whether the CGI Mountain score is larger than zero.

##### Simple repeats

We annotated the simple repeat regions in the genome according to RepeatMasker (http://www.repeatmasker.org), which defines the interspersed and low-complexity repeats in hg19. We use a dummy variable to indicate whether a genomic site is in a region masked as simple repeats.

##### DNase I peaks

We defined DNase I peaks according to the DNAse I annotation from the Roadmap Epigenomics project [Roadmap Epigenomics Consortium, 2015]. We use the score from HoneyBadger2 to indicate the DNase I signal strength for regions with a DNase I peak (http://www.broadinstitute.org/~meuleman/reg2map/). The regions not annotated in the HoneyBadger2 were masked as “no peak” regions in our analysis.

##### PhyloP score

The conservation score phyloP (phylogenetic *p*-values) is part of the PHAST package (http://compgen.bscb.cornell.edu/phast/). We used the score from the multiple alignments of 99 vertebrate genomes to the human genome [Pollard et al., 2010]. We use the version of phyloP100way which covers 99.8% of the subtracted regions.

##### Replication timing

We adjust the replication timing data from Chen et al. [2010] to the hg19 assembly. The replication timing value ranges from 0 to 1, indicating the earlier to later stages in replication process. Annotation of replication timing covers 91.2% of the subtracted regions.

##### Gene expression

We define the gene expression level according to TCGA RNAseq expression data. Expression data was log_2_(*x* + 1) transformed. For each cancer type, the median expression was calculated for all genes. If multiple annotations of a gene exist, the longest annotation is used. For overlapping genes, the expression is a cumulative sum.

As in Fredriksson et al. [2014], we collapse colon (COAD) and rectal carcinoma (READ) to a joint cancer type CRC by averaging across the expression values.

#### 4.1.2 Preparation of the analytical data table

In order to facilitate the model fitting procedure, we summarized the genomic data into count tables. In Poisson count models, each row in the count table represents a pre-defined region. For continuous explanatory variables, such as replication timing, the annotations are averaged over the region. For categorical explanatory variables, the annotations are transformed to the percentage for different levels of the variables, e.g, the binary explanatory variable indicating whether the site is a (G:C) base pair or not is transformed to GC content of the window. In site-specific regression models, we discretized continuous variables into bins according to quartiles or quintiles. Each row in the count table represents the counts of mutations under a certain combination of levels of all the explanatory variables. As we are summarizing the whole genome in the count table, we expect to see all the combinations of the levels for all the explanatory variables. Thus, the sizes of the count tables grow significantly when adding new explanatory variables. The generation of the count tables also takes much longer time with more explanatory variables. Because of the space and time consumption, a large count table including all the explanatory variables is computationally infeasible. We implemented the forward model selection procedure to avoid many huge count tables. For each iteration in the forward model selection procedure, we made new count tables only for the selected sets of explanatory variables from previous step and one new candidate explanatory variable. We then added the best candidate in the explanatory variable set and repeated for the next step. We used 2% of the whole genome in the model selection and made a final count table with the remaining 98% sites only for the important explanatory variables that were determined from the model selection procedure. The generation of the count table for the final model takes 6,000 CPU hours on our cluster (2.5GHz CPUs).

### 4.2 Multinomial regression model

#### 4.2.1 Estimation

The observations in the regression model are indexed by the genomic position *i*(1 ≤ *i* ≤ 2.56 · 10^9^) for all positions on chr1 to chr22 after excluding problematic regions and the samples *sam*(1 ≤ *sam* ≤ 505), so the total number of observations is 1.3 · 10^12^. Starting from Model 4, sample specific intercepts for the six mutation types sum up to 6 · 505 = 3030 parameters. The regression coefficients for explanatory variables are indexed by the 14 cancer types. In Model 5 each cancer type has 4 · 4 · 6 = 96 parameters for the neighboring context and the mutation type (4 for the left neighboring site, 4 for the right neighboring site and 6 for the mutations), so the total number of parameters in this model is 6 · 505 + 14 · 96 = 4374. From Model 6 to Model 8 we use 1 parameter for each of the continuous explanatory variables phyloP, replication timing and expression level. We also add 1 dummy variable for expression level indicating whether the given site is potentially expressed or not. In Model 6 we have 4374 + 14 · 6 = 4458 parameters. In Model 7 we have 4458 + 14 · 6 = 4542 parameters. In the final model we have 6 · 505 + 14 · 96 + 14 · (3 + 1) · 6 = 4710 parameters in total. To reduce memory usage and computation time, the parameters are estimated in three separate binary logistic regression models, but the variance-covariance matrix of the parameters is estimated for the multinomial model [Begg and Gray, 1984]. Estimation is conducted in R [R Core Team, 2014] using the function glm4 from the contributed package MatrixModels [Bates and Maechler, 2015b] and the estimation of the variance-covariance matrix is implemented using the package Matrix [Bates and Maechler, 2015a].

#### 4.2.2 Dirichlet prior and pseudo counts

If a model with many explanatory variables and interaction terms among them is estimated (e. g. sampleID × neighbors × strong), it can easily occur that for a certain combination of levels of categorical variables, there have been no mutations of a certain type observed. This case is especially likely, if the sampleID is involved. This causes numerical problems in the maximum likelihood estimation [Agresti, 2002].

To solve this problem while still obtaining a positive mutation probability, we added pseudo counts to the observed mutation counts. This is equivalent to specifying a Dirichlet prior for the multinomial model [Durbin et al., 1998]. We obtain the pseudo counts from the mutation counts of the corresponding cancer type for the exact same combination of categorical variables. The mutation counts from the sample and from the cancer type are equally weighted.

#### 4.2.3 McFadden’s pseudo *R*^2^

To assess the fit of a model, we report McFadden’s pseudo *R*^2^

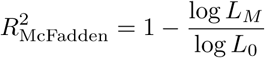

where *L_M_* is the likelihood of the model under investigation and *L*_0_ is the likelihood of a model with no predictors [McFadden, 1974]. To measure the improvement of a model in comparison with the binomial model where there is no distinction between the mutation types, we use the same mutation probability for each mutation type in the model without predictors.

### 4.3 Forward variable selection

To speed up data preparation and estimation, variable selection is conducted on a subset of approximately 2% of the genome. It is constructed by randomly selecting 60,000 windows of size 1 kb. The cancer types kidney chromophobe, low-grade glioma, prostate adenocarcinoma and thyroid carcinoma have very low mutation counts, so they are disregarded during variable selection. We use cross-validation for forward variable selection. The improvement of the fit along the model selection procedure are measured with McFadden’s pseudo *R*^2^ and deviance loss.

Starting with Model 5,

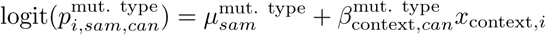

additional terms of the form

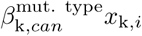

with explanatory variable k ∊ {phyloP, replication timing, genomic segment, expression, GC content, DNase1, simple repeats, CpG island}, are selected following the forward selection scheme.

#### 4.3.1 Cross validation

An assessment of the fit of a model can be obtained by cross validation. Five-fold cross validation is used to select the annotation with the highest explanatory power. To this end, the 1 kb windows of the variable selection subset are divided randomly distributed among 5 sets. In turn, 4 of them are joined as the training set, on which the multinomial model is estimated, and the remaining set is used as the validation set to estimate the loss function.

The deviance loss function for the observed site *i* in sample *sam* is defined as

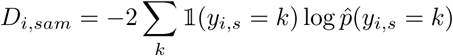

where *k* denotes all possible mutation events at site *i, y_i_*_,*sam*_ the observed event at site *i* in sample *sam* and 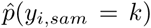 the probability of event *k* estimated under the multinomial regression model [Hastie et al., 2001]. Thus, it measures the prediction accuracy of the multinomial regression model. The total deviance loss of a model is

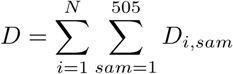

with *N* is the number of genomic positions.

## Appendix: Forward model selection results

In this appendix, we provide the details for the forward model selection procedure.

**Step 1**

**Table 1:**
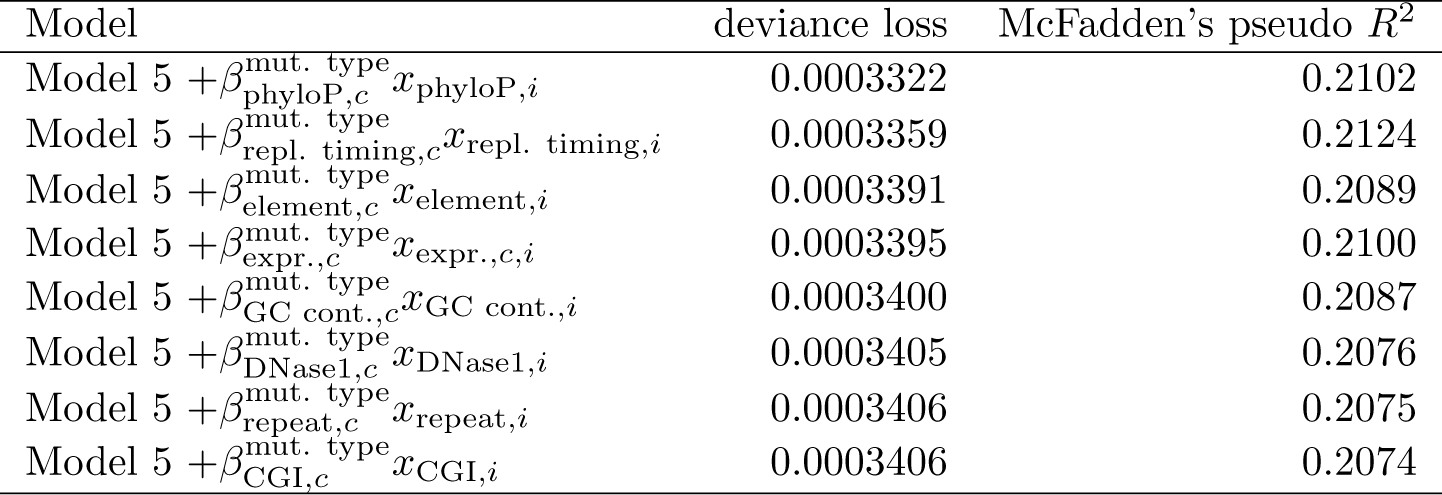
Deviance loss and McFadden’s pseudo *R*^2^ for each of the models tested in step 1 to obtain model 6.

**Step 2**

**Table 2:**
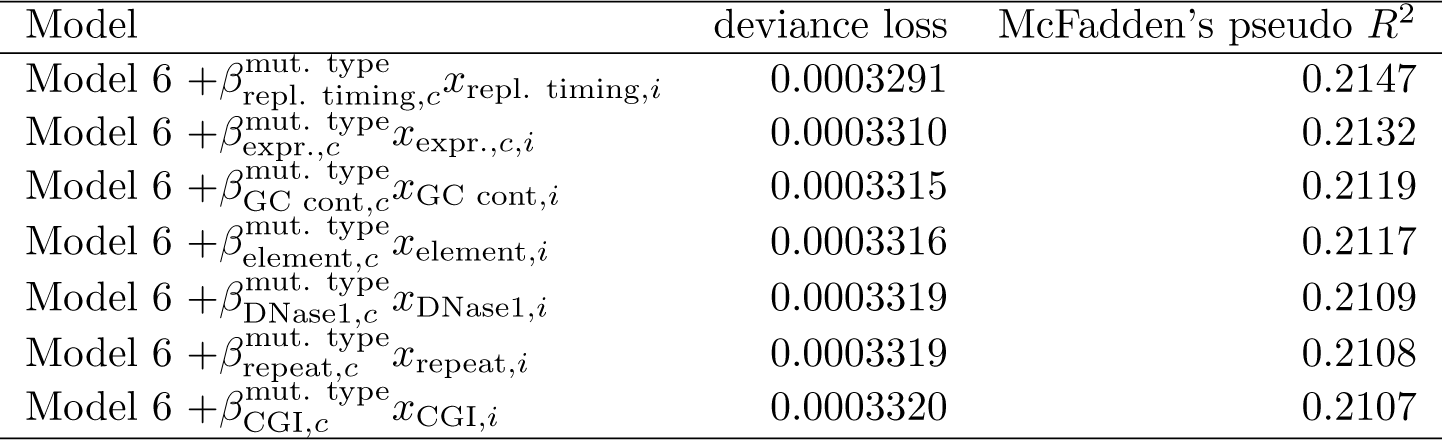
Deviance loss and McFadden’s pseudo *R*^2^ for each of the models tested in step 2 to obtain model 7.

**Step 3**

**Table 3:**
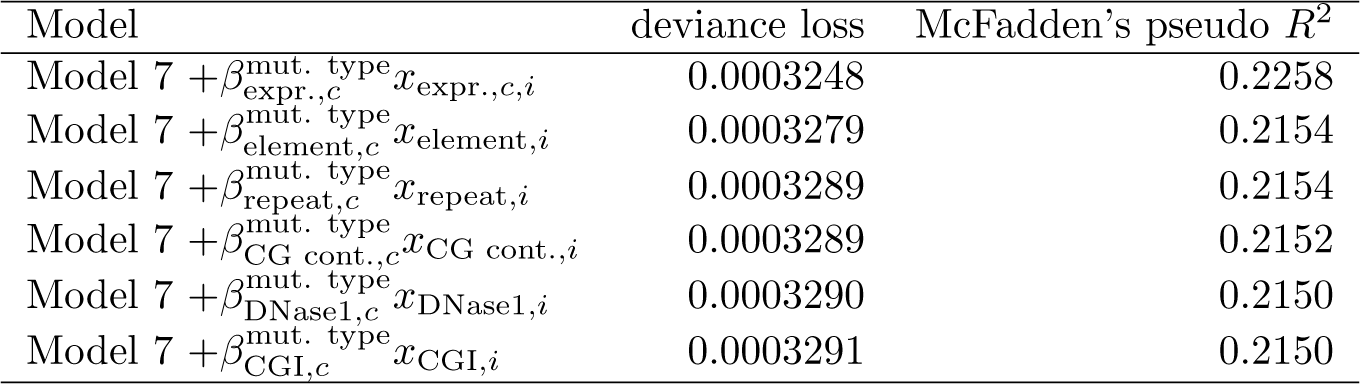
Deviance loss and McFadden’s pseudo *R*^2^ for each of the models tested in step 3 to obtain model 8.

## List of abbreviations

TCR: transcription-coupled repair
NER: Nucleotide excision repair

## Declarations

### Ethics approval and consent to participate

Not applicable.

### Consent for publication

Not applicable.

### Availability of data and material

The sources of the datasets used in this article are well described in Section 4.1. The analysis and final result of our model are available from the corresponding authors.

### Competing interests

Not applicable.

### Funding

Not applicable.

### Authors’ contributions

JB, QG, MJ, SB, AH and JSP designed the logistic regression framework. JB, QG, MJ, SB, MMN, HH and JSP collected the datasets for explanatory variables and mutation calls. JB, QG and MJ analyzed the datasets. All authors contributed to the interpretation of the results. JB and QG drafted the manuscript. AH and JSP revised the manuscript critically for important updates. All authors read and approved the final manuscript.

## Acknowledgements

Not applicable.

